# A new and widespread group of fish apicomplexan parasites

**DOI:** 10.1101/2024.01.10.575052

**Authors:** Anthony M Bonacolta, Joana Krause, Nico J Smit, Paul Sikkel, Javier del Campo

## Abstract

Apicomplexans are obligate intracellular parasites which have evolved from a free-living, phototrophic ancestor. Well-known representatives include *Plasmodium*, the causative agent of malaria, and the gregarines which infect a vast number of marine invertebrates. Apicomplexans have been reported from marine environmental samples in high numbers^1^, with several clades of Apicomplexan-Related Lineages (ARLs) having been described from environmental sequencing data (SSU rRNA metabarcoding)^2^. The most notable of these being the corallicolids (previously known as ARL-V) which possess chlorophyll-biosynthesis genes in their relic chloroplast (apicoplast) and are geographically wide-spread and abundant symbionts of anthozoans^3^. *Ophioblennius macclurei*, the redlip blenny, along with other tropical reef fishes, is known to be infected by *Haemogregarina*-like and *Haemohormidium*-like parasites^4^; however previous phylogenetic work using the 18S rRNA genes shows these parasites to be distinct from where they are classified morphologically^5,6^ and instead related to the Coccidia. Hybrid genomic sequencing incorporating Oxford Nanopore Technologies long reads and Illumina short reads of apicomplexan-infected *O. macclurei* blood was carried out to retrieve more genetic information on these parasites. Using this approach, we were able to recover the entire rRNA operon of this apicomplexan parasite along with the complete mitochondrion and apicoplast genomes. Phylogenetic analyses using this new genomic information consistently place these fish-infecting apicomplexans, hereby informally named ichthyocolids, sister to the corallicolids within Coccidia. The apicoplast genome did not contain chlorophyll biosynthesis genes, providing evidence for another independent loss of this pathway within Apicomplexa. Based on the 16S rRNA gene found in the apicoplast, this group corresponds to the previously described ARL-VI. Screening of fish microbiome studies using the plastid 16S rRNA gene shows these parasites to be geographically and taxonomically widespread in fish species across the globe with potentially major implications for commercial fisheries and oceanic food webs.

## RESULTS & DISCUSSION

*Haemogregarine*-like and *Haemohormidium*-like parasites of tropical fishes were first classified based on morphological identification of life stages observed in erythrocytes^4,7^, however further phylogenetic work using the 18S rRNA gene placed them in a previously uncharacterized clade within Coccidia^5,6^. The clade Coccidia includes wide-spread parasites of mammals, birds, and other animals, as well as recently described symbionts of corals, commonly known as the corallicolids^3^. Parasites morphologically identified as *Haemohormidium*-like were more prevalent in Caribbean damselfish^8^, while those identified as *Haemogregarine*-like were more prevalent in blennies^6^. To better characterize these fish-infecting apicomplexans, a hybrid metagenomic assembly incorporating Oxford Nanopore Technologies long reads and Illumina short was constructed from the blood of the redlip blenny, *Ophioblennius macclurei,* with an acute infection of a *Haemogregarine*-like parasite (confirmed by Giemsa staining; Figure 1A). By analyzing the parasite reads in these data, we were able to recover the complete rRNA operon, as well as assemble organelle genomes for this apicomplexan (henceforth referred to as an ichthyocolid, meaning ‘fish-dweller’ from Greek-derived Latin *ichthys* combined with the suffix –*cola*, derived from the Latin *incola,* following the naming conventions used in Kwong et al. 2019 for the corallicolids). In addition, we screened published fish microbiome studies for the presence of this parasite across fish taxonomic and geographic diversity.

**Figure 1.**
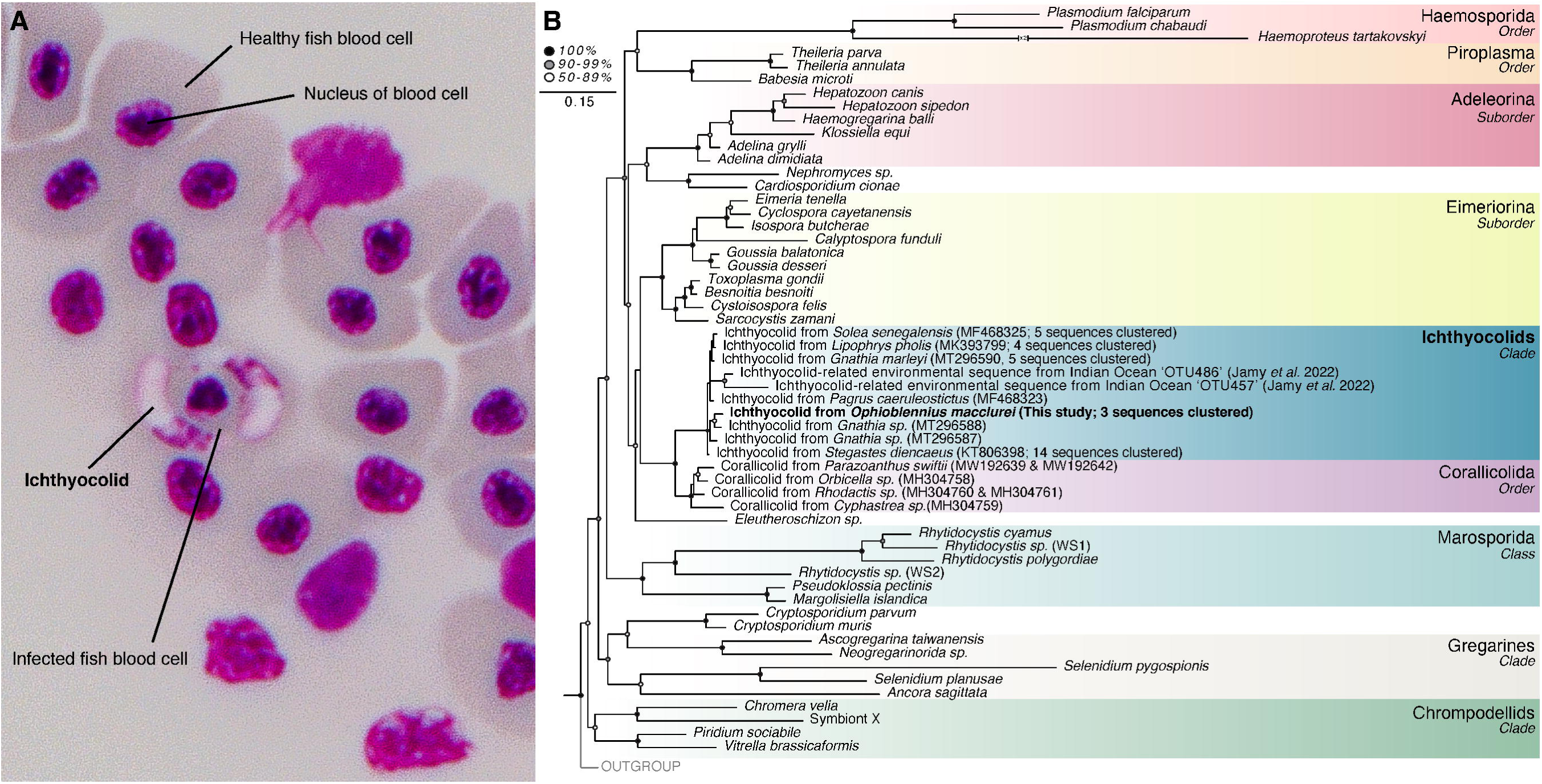
Giemsa-stained *Ophioblennius macclurei* blood & rRNA operon tree of Apicomplexa. (A) *Haemogregarine*-like apicomplexan parasites within the blood cells of *O. macclurei* stained with Giemsa. (B) Maximum-likelihood rRNA operon tree of Apicomplexa. This tree was constructed using the 18S and 28S subunits of the rRNA operon using 5 074 nucleic acid sites with RAxML GTR substitution matrix with GAMMA model of rate heterogeneity and 1 000 bootstrap replicates.

We built a phylogenetic tree of the rRNA operon incorporating the recovered 18S and 28S of our ichthyocolid with the 18S of the fish apicomplexans recovered in Sikkel et al. 2018^8^, Hayes et al 2019^6^, and Renoux 2017^5^, the 18S and 28S of coccidians recovered from Jamy et al. 2022^9^, the *Ophioblennius macclurei* sequence noted above (KY940307), and the 18S-28S of other representative apicomplexans. We find that our parasite forms a strongly supported clade (100% bootstrap) with other fish parasites sister to the corallicolids, which we term the ichthyocolids (Figure 1B). Previously described *Haemogregarine*-like and *Haemohormidium*-like parasites of marine fishes all fall within this clade, despite slight differences in their morphology. There appears to be a level of host-specificity within the ichthyocolid clade as well, with all the *Haemohormidium*-like ichthyocolid sequences originating from damselfish (*Stegastes spp.)* clustering together and our *Haemogregarine*-like ichthyocolid recovered from this study clustering with the *Ophioblennius macclurei* sequence (Figure 1B). Furthermore, we found two 18S-28S OTUs from Jamy et al 2022^9^ previously assigned to the corallicolids actually clustered sister to the ichthyocolid clade. Both environmental sequences were recovered from a 0.2-3 μm size-selected sample from 800 m deep in the Indian Ocean. Although we cannot confidently classify these sequences as ichthyocolids, we hypothesize these ichthyocolid-related OTUs to likely come from cyst life-stages found within the water column or potentially from inside planktonic vectors of the parasite.

Having established our parasite as sister to the corallicolids, we conducted a thorough exploration of our hybrid metagenomic assembly to extract further genetic information on the ichthyocolids. This investigation successfully unveiled the 6.3 kb linear mitochondrion genome (Figure 2A). The size of the ichthyocolid mitochondrion is consistent with the extremely reduced mitochondrion genomes found within Apicomplexa, which are known to be among the smallest found in nature^10^. It is the same size as the corallicolid mitochondrial genome^11^. Like most other apicomplexans, the ichthyocolid mitochondrion genome encodes only three genes: *cox1*, *cox3*, and *apocytochrome b* (*cob*); with *cox1* and *cob* being fused together (Figure 2A). The apicomplexan related algae *Vitrella spp.* also possesses a mitochondrial genome with a fused *cox1* and *cob*^12^. Like the rRNA operon phylogeny, the mitochondrial phylogeny using a concatenated alignment of the *cox1*, *cox3*, and *cob* genes strongly supports (100% bootstrap) ichthyocolids as sister to the corallicolids (Figure 2B). This clade is within Coccidia and is sister to the Eimeriorina suborder which includes *Toxoplasma gondii* and *Eimeria tenella* (Figure 2B).

**Figure 2.**
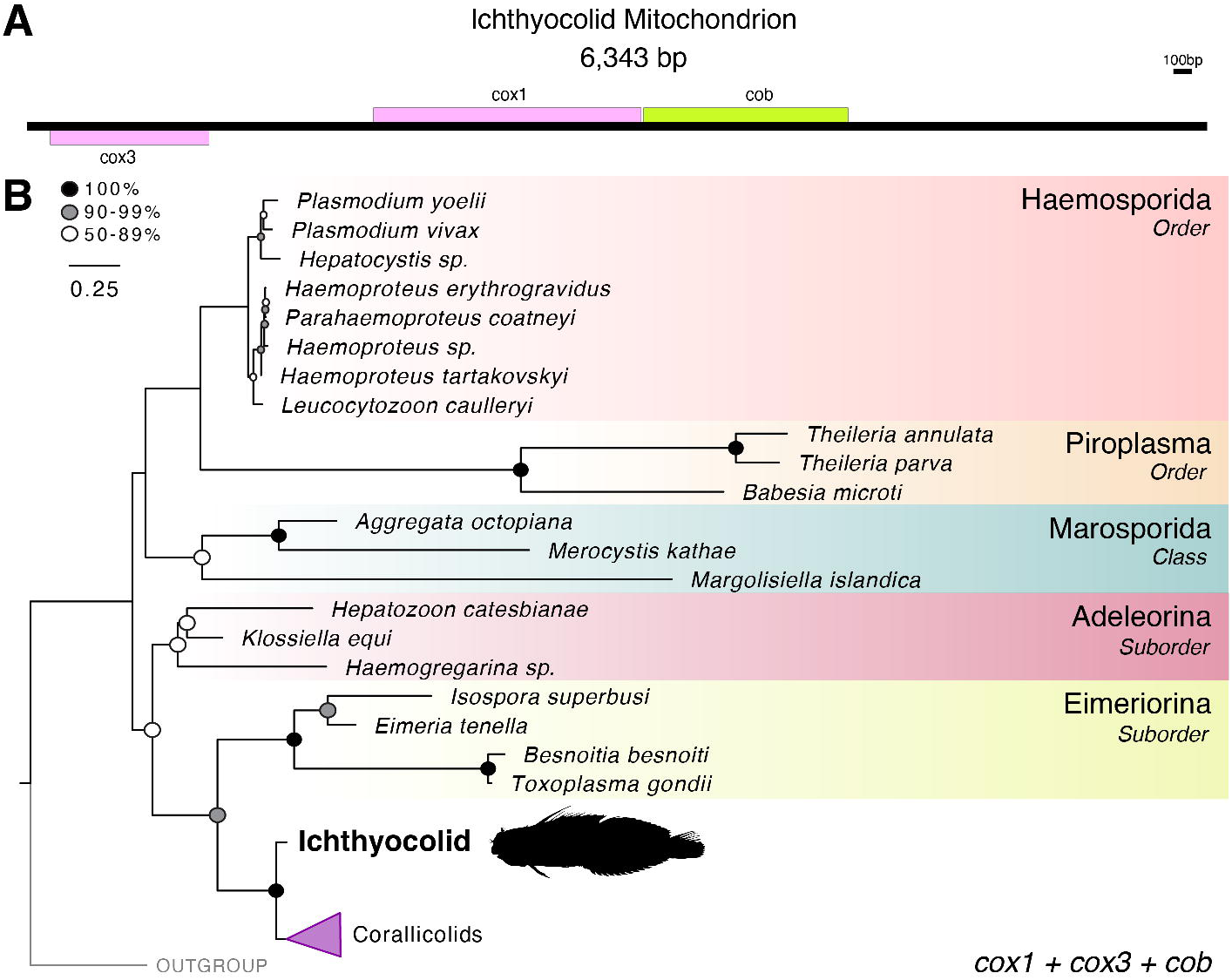
Ichthyocolid mitochondrial genome & phylogenetics. (A) Visualization of the linear mitochondrial genome of the ichthyocolids. (B) Maximum-likelihood tree of apicomplexan mitochondria based on *cox1*, *cox3*, and *cob* using 1 322 amino acid sites using RAxML MTZOA substitution matrix with GAMMA model of rate heterogeneity and 1 000 bootstrap replicates. Major clades of Apicomplexa and related lineages are color-coded. Stramenopiles were used as the outgroup.

After recovery of the mitochondrion genome, we sought to assemble the genome of the other primary organelle found within Apicomplexa, the apicoplast. This vestigial plastid, which is homologous to chloroplasts, is responsible for generating essential molecules which the symbiont cannot scavenge from its host(s)^13^. Using GetOrganelle^14^ we were able to assemble a complete, circularized 39 kb apicoplast genome from our parasite (Figure 3A), which is smaller than that of the corallicolids (46 kb)^3^ but larger than that of *Plasmodium sp.* (35 kb)^15^ and *Eimeria tenella* (35 kb)^16^. Apicoplast phylogeny constructed from a concatenated alignment of 31 apicoplast genes strongly supports (100% bootstrap) the ichthyocolid’s plastid as sister to that of the corallicolids, once again forming a distinct well-supported clade within Coccidia (Figure 3B). This relationship is also supported by the revelation that the parasite’s plastid gene-content is highly similar to that of the corallicolids, with the ichthyocolid only missing the chlorophyll biosynthesis pathway genes (*acsF*, *chlB*, *chlL*, *chlN*), *rps17*, *orfB*, *orfF*, and *orfE* (Figure 1C). Additionally, along with our parasite, the corallicolids possess the only other apicoplast genome encoding the 5S rRNA. Like all other apicoplasts^17^, ancestor chloroplast-associated genes *rps14*, *rpl3*, *rpl27*, and *rpoA* are not present (Figure 3C). Although chlorophyll biosynthesis genes were not found in the apicoplast of the ichthyocolid, this does not rule out that these genes may have been transferred to the nucleus via endosymbiotic gene transfer and have not yet been fully lost.

**Figure 3.**
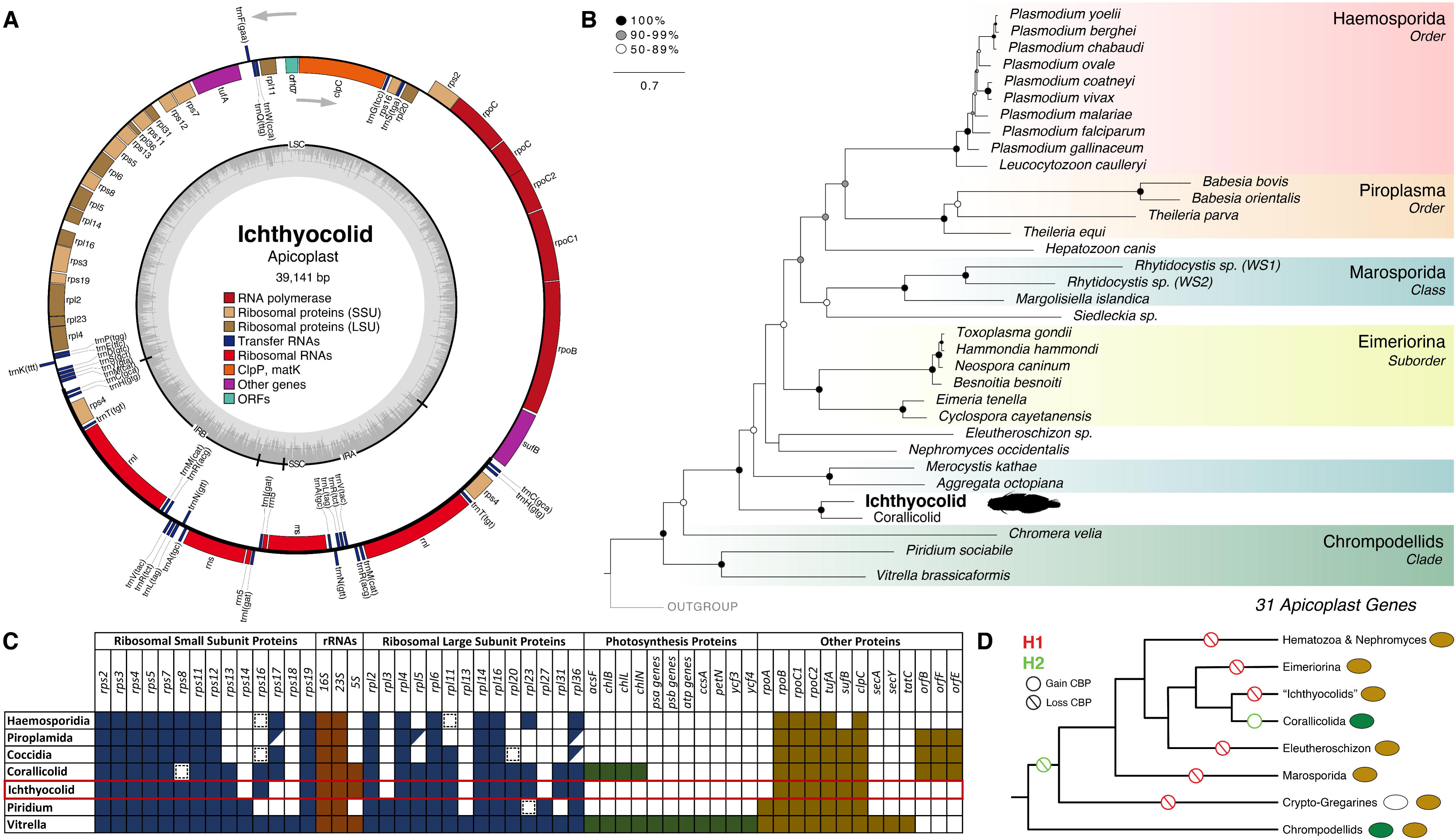
Ichthyocolid apicoplast genome & phylogenetics. (A) Visualization of the apicoplast genome of the ichthyocolids. (B) Maximum-likelihood tree of apicomplexans based on 31 plastid-encoded genes and 10 086 amino acid sites using RAxML MTZOA substitution matrix with GAMMA model of rate heterogeneity and 1 000 bootstrap replicates. Major clades of Apicomplexa and related lineages are color-coded. Rhodophyte plastids were used as the outgroup. (C) Plastid gene content of apicomplexans and related lineages. Variable genes are represented with a slash, while remnant genes are represented by a dashed box. (D) Loss of the chlorophyll biosynthesis pathway (CBP) through apicomplexan evolution based on the most recent phylogenomic topology. Brown ovals indicate lineages with plastids without the chlorophyll biosynthesis pathway, green ovals indicate ones with it, and empty ovals represent a complete loss of the plastid in that lineage.

The absence of the chlorophyll biosynthesis pathway in the ichthyocolids, despite being the closest known relative to the corallicolids is relevant as it likely represents another independent loss of the pathway within Apicomplexa and suggests that the corallicolids may be unique in their retention of it. This is surprising given the predilection of this lineage towards the convergent loss of photosynthesis and the associated chlorophyll biosynthesis genes, which precluded the multiple independent origins of apicomplexan parasites^17^. While a parsimonious interpretation of evolution (shown by H2 in figure 3D) would presume one gain of the chlorophyll biosynthesis pathway in the corallicolids after an initial loss of it in the last common ancestor of the Apicomplexa, the more plausible scenario is the one (shown by H1 in figure 3D) where the chlorophyll biosynthesis pathway is repeatedly lost throughout apicomplexan evolution. Our results contribute to this hypothesis with the 6th apparent loss of the pathway (based on the latest phylogenomic analysis^18^) within apicomplexans occurring in the ichthyocolids (Figure 3D). The reduction of the apicoplast genome through evolution is marked by the loss of photosystem genes from the common ancestor with chromerids, followed by independent, repeated loss of the chlorophyll biosynthesis pathway^3^. Thus, the last common ancestor of the corallicolids and ichthyocolids likely possessed chlorophyll biosynthesis genes.

As we were able to recover the complete apicoplast of our ichthyocolid, we extracted the 16S rRNA gene from this organelle to correlate its phylogeny with previously identified apicomplexan-related lineages (ARLs) from environmental barcoding data. Janouškovec et al. 2012 identified numerous apicomplexan-related lineages (ARLs) from environmental barcoding data using plastid 16S (SSU) rRNA sequences, revealing a vast underexplored diversity of alveolates in the marine environment^2^. Later work on one of these lineages, ARL-V, led to the full characterization of this group as the corallicolids^3,19^. Incorporating the apicoplast SSU rRNA we recovered from our ichthyocolid into the phylogenetic framework presented in Janouškovec et al. 2012^2^ & 2019^20^, as well as with additional corallicolid apicoplast SSUs from Kwong et al. 2021^19^, we were able to identify our parasite as a member of ARL-VI, henceforth referring to other members of this clade as ichthyocolids as well (Figure 4A). The other members of this clade were isolated from fish specimens, indicating a potential exclusive association of these blood parasites with fish as the primary host.

**Figure 4.**
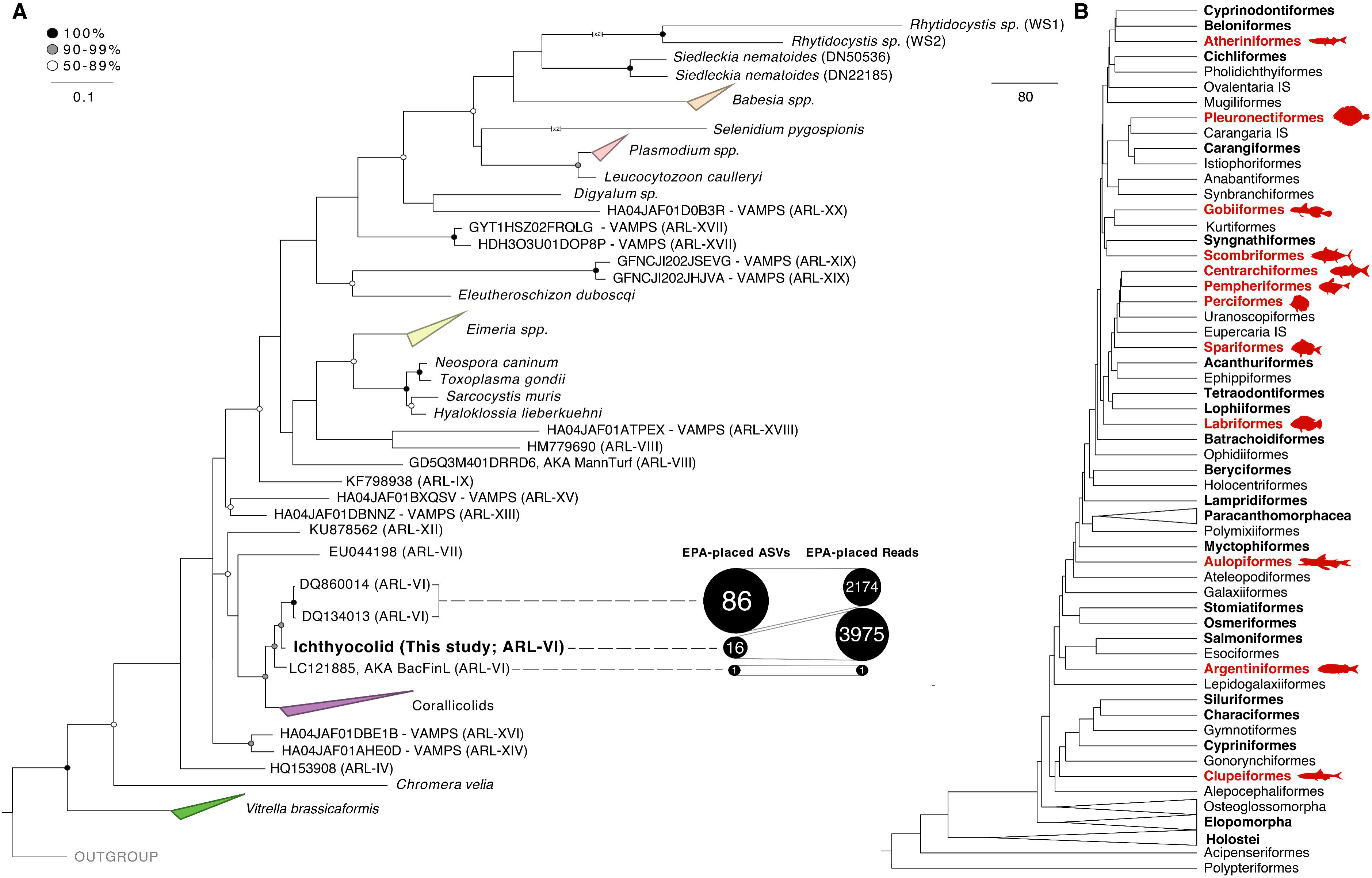
Apicomplexan-related lineages (ARLs) based on Plastid 16S rRNA & likely ichthyocolid susceptible fish orders. (A) Maximum-likelihood tree of Apicomplexa using plastid 16S rRNA (SSU) based on 1 398 nucleic acid sites using RAxML GTR substitution matrix with GAMMA model of rate heterogeneity and 1 000 bootstrap replicates. Rhodophyte plastid SSUs were used as the outgroup. The amount of amplicon sequence variants (ASVs) recovered from fish microbiome studies which either cluster with or are EPA placed within certain isolated ichthyocolids is shown in bubbles along with the total number of reads that correspond to these ASVs. (B) Cladogram of Teleost orders showing which clades were included in the microbiome datasets screened (bold) and which ones had ichthyocolid reads recovered from their microbiome (bold red text and silhouette).

To uncover the potential diversity of the newly characterized ichthyocolids we took advantage of the ample publicly available fish microbiome datasets (16S rRNA + metagenomic) covering marine and freshwater teleost (bony fishes) as well as elasmobranchs (cartilaginous fishes) and screened a comprehensive collection of 14 datasets for the apicoplast SSU rRNA. Unfiltered datasets were first checked via BLASTn for sequences similar to the ichthyocolid apicoplast SSU rRNA query (90% similarity threshold), then all the recovered amplicon sequence variants (ASVs) were mapped using the evolutionary placement algorithm in RAxML to an apicoplast SSU rRNA tree to confirm their identity (Figure 4A). Of the ASVs clustering with the ichthyocolids, 86 clustered closely with the isolates from oceanic anchovies (DQ860014 and DQ184013), while 16 clustered closely with the *O. macclurei* ichthyocolid described in this study (Figure 4A). One ASV clustered with an ichthyocolid found within the flathead grey mullet, *Mugil cephalus*, from a Mediterranean estuary (LC121856; Figure 4A). Ichthyocolids do not appear in any of the screened elasmobranch microbiomes (n_publications_ = 3; n_orders_= 6). Neither do they appear in freshwater teleost microbiomes (n_publications_ = 3; n_orders_= 8). However, ichthyocolids appear to be widespread in marine teleost microbiomes with ichthyocolid-like reads appearing in 37 fish species, encompassing 23 different families and 11 orders (Supplementary Table 1; Figure 4B). In total ∼36% of the fish orders screened were infected with likely ichthyocolids, including the highly specious *Perciformes*. Infected teleosts also include those important to major fisheries such as tuna (*Scombriformes*) and anchovies (*Clupeiformes*). These fish samples represent a wide variety of tissues (skin, blood, gills, & internal organs), vast geographic distribution (Pacific Ocean to Atlantic Ocean), and habitat types (open ocean, deep sea, coral reefs, kelp forests, tidepools, etc.; Supplementary Table 1). These results support ichthyocolids as being a globally distributed, general teleost parasite with under-sampled diversity. Additionally, recent work shows that fish infected with this parasite have high leukocyte counts^21^ and are likely immune compromised suggesting further exploration of the ecological impact of the fish parasites is needed.

Potential vectors include ectoparasites such as leeches and gnathiids, as each have been documented as vectors of blood-borne apicomplexans before^22–24^. Gnathiid isopods represent the most likely vectors for these parasites as ichthyocolids 18S rRNA sequence reads have been recovered from them directly and is an avenue of future exploration^25^.

Using genetic information retrieved from the hybrid-read assembled metagenome of parasite-infected fish blood and phylogenetic analyses we have been able to classify the *Haemogregarine*-like parasite of *O. macclurei* as a member of a new clade of fish-infecting apicomplexans we refer to as the ichthyocolids. Through organelle multi-gene trees and rRNA gene trees, we found this clade to be sister to the corallicolids and composed of coccidians predominately found as blood parasites of fish species across the globe. Using the SSU of the ichthyocolid plastid, we show that this new clade corresponds to the previously described ARL-VI. The plastid SSU was also used to assess the extent and potential distribution of this parasite in the world’s oceans by mining previously published fish microbiome datasets, confirming its presence in a variety of commercially important fish species, from tuna to anchovies and flatfishes. The ichthyocolids plastid does not contain chlorophyll biosynthesis genes, like the plastids of the corallicolids, and thus represent another independent loss of this pathway within Apicomplexa. Future work should further define the diversity of this group and its interesting evolutionary history. The other life stages of this parasite are also of interest and more information on their transmission through their likely invertebrate vectors, gnathiid isopods, is necessary. Critically, the ecological and commercial impacts of this parasite need to be further assessed given its wide-distribution and potential impact on vital fisheries and oceanic food webs.

## Acknowledgments

We would like to acknowledge Bradley Weiler for assisting with sample collection in the middle of the COVID-19 pandemic. We would also like to thank those who shared fish microbiome data with us, specifically Dr. Jeremiah Minich for his 101 Marine Fish Microbiome dataset^26^. This study has been supported by project PID2020-118836GA-I00 financed by MCIN/AEI/10.13039/501100011033, by project 2021 SCR 00420 financed by Departament de Recerca i Universitats de la Generalitat de Catalunya, OCE 2023420 from the National Science Foundation, and startup funds from the University of Miami, Rosenstiel School of Marine, Atmospheric and Earth Sciences.

## Author Contributions

Conceptualization, J.d.C., P.S.; Methodology, J.d.C., P.S., N.S., A.M.B; Formal Analysis, A.M.B, J.K.; Investigation, J.d.C., P.S., N.S., A.M.B.; Resources, J.d.C. P.S.; Data Curation, A.M.B. and J.d.C.; Writing – Original Draft, A.M.B; Writing – Review & Editing, A.M.B, P.S., N.S., J.K., J.d.C.; Visualization, A.M.B., J.K.; Supervision, J.d.C.; Project Administration, J.d.C.; Funding Acquisition, J.d.C.

## Declaration of Interests

The authors declare no conflicts of interest.

## STAR Methods

### Method Details

#### Sample Collection & Confirmation of Parasite Prescence

In October 2020, *O. macclurei* were caught off reefs in Curacao and transported, alive, by boat to Summerland Key, Florida, USA. Blood was extracted from the caudal artery of these fish using a sterile 26-gauge needle and syringe. Blood smears were made by allowing a single drop of blood to air-dry for 10 minutes before fixation with absolute methanol. These smears were later stained using a modified Giemsa stain solution then examined for parasite presence^7^. The remaining blood was immediately spun down at 600 x g for 5 minutes at 4C. The pellet was preserved in 80% EtOH and frozen at -20C. DNA was extracted using the QIAGEN DNeasy Blood and Tissue Kit (QIAGEN, Hilden, Germany). Extracted DNA was cleaned of impurities using AMPURE MagBeads (Beckman Coulter, Indianapolis, USA) at 1.8x ratio and extraction quality was assessed using a Qubit 4 Fluorometer (Thermo Fisher Scientific, Waltham, USA) and a Nanodrop Spectrophotometer (Thermo Fisher Scientific, Waltham, USA).

#### Sequencing & Metagenome Assembly

High-quality DNA was used as input to generate nanopore sequencing libraries using the Nanopore Genomic DNA by Ligation protocol (Version: GDE_9063_v109_revAB_14Aug2019; Kit: SQK-LSK109). Sequencing was conducted using a MINion FLO-MIN106 Flowcell. Basecalling was performed using Guppy-CPU on University of Miami’s Pegasus Super Computer. 381 370 raw Nanopore reads were corrected and scrubbed using Miniscrub2 v0.4^27^. Quality control of the long reads was conducted using LongQC v1.2.0c^28^. For short reads, 100 M PE150 reads were generated using the Illumina Novoseq 6000 at John P. Hussman Institute for Human Genomics (University of Miami) from the same DNA input. Fastp v0.23.4was used for adapter trimming and quality control^29^. The Nanopore and Illumina reads were used together as input to generate a hybrid metagenomic assembly of the fish and parasite reads using Opera-MS v0.8.3^30^.

#### rRNA Phylogenetic Analysis

To recover the rRNA operon, barrnap v0.9^31^ was used to extract the SSUs and LSUs from the metagenomic assembly. BLASTn v2.13.0 was used to identify the parasite SSU and LSU from the extracted set^32^. As both apicomplexan-derived subunits were derived from the same contig in the metagenomic assembly, the entire sequence spanning from the SSU to the LSU was extracted as the rRNA operon. The 18S-28S rRNA operon phylogenetic tree of Apicomplexa was constructed by combining the ichthyocolid 18S-28S from this study with the likely ichthyocolids from Sikkel *et al.* 2018^8^, coccidian 18S-28S OTUs from Jamy *et al.* 2022^9^, and other representative apicomplexan sequences. Sequences longer than 500 bp and shorter than 15 000 bp were sorted and clustered at 98%, then aligned using MAFFT v7.480. The alignment was then trimmed using Trimal v1.4.rev15^33^ with a gap threshold of 30%, similarity threshold of 0.001, and a minimum conservation of 30%. A maximum-likelihood tree was then constructed using RAxML v8.2.12 with the General Time Reversible model with a gamma-distributed rate variation (GTR+CAT) and 1 000 bootstrap replicates^34^.

#### Organelle Genome Assemblies and Phylogenetic Analyses

Contigs from the metagenomic assembly were binned using metabat2, MaxBin2, and CONCOCT using a Metawrap wrapper script^35–38^. Metagenomic bins were then taxonomically classified using CAT/BAT v5.2.3 with the NCBI database^39^. Bins identified as apicomplexans were joined and BLASTn was run to identify the contigs^32^. A full mitochondrion genome and a preliminary apicoplast genome were identified and these contigs were used as seed inputs for GetOrganelle v1.7.7^14^ to generate a complete and circularized apicoplast genome. MFANNOT v1.36^40^ was used to annotate both the apicoplast and mitochondrion genomes using genetic code 4 as done previously with apicomplexan organelles^3^. OrganellarGenomeDRAW v1.3.1 was used to visualize the organelle genomes^41^.

For the apicoplast multi-gene tree, previously published proteomic datasets (Genebank in supp excel) were combined with the ichthyocolid data^17,19^. Custom python scripts were then used to extract genes from the assemblies and compile into shared gene sets. MUSCLE v3.8.1551 was used to generate alignments for each gene set^42^. Gene set alignments were then concatenated using Concatenator v0.2.1^43^. A final concatenated alignment of 31 plastid-encoded genes was used to construct a phylogenomic tree using RAxML with the MTZOA substitution matrix and GAMMA model of rate heterogeneity and 1 000 bootstrap replicates ^34^. For the mitochondrion multi-gene tree, MITOS v1.1.1 was run to translate our mitochondrial genome then the cox1, cox3, and cob genes were combined with other apicomplexan mitochondrial sequences derived from genbank (excel supp)^44^. Dinoflagellates were used as the outgroup as done for previous apicomplexan mitochondrial tree reconstructions as the chrompodellids show extreme divergences in their mitochondrial genomes. Alignment, concatenation, and phylogenomic tree construction was done as before for the apicoplast genes.

Like the ichthyocolid rRNA operon above, the apicoplast SSU was identified using barrnap^31^. It was then combined with the datasets from Janouškovec et al 2012, Janouškovec et al 2019, & Kwong et al 2020^2,19,20^. Sequences longer than 300 bp and shorter than 2 500 were sorted and clustered at 99.5%, then aligned using MAFFT^45^. The alignment was then trimmed using Trimal with a gap threshold of 30%, similarity threshold of 0.001, and a minimum conservation of 30%^33^. A maximum-likelihood tree was then constructed using RAxML (GTR+CAT model) with 1 000 bootstrap replicates.

#### Fish Microbiome Screening

Unfiltered ASV sequences from fish 16S rRNA metabarcoding datasets as well as raw reads from shotgun metagenomic datasets were screened for the presence of the ichthyocolid apicoplast SSU. The 14 microbiome studies encompassing 46 different orders of fish included in this analysis can be found in Supplementary Table 2. First a BlastDB was constructed using the sequences from each study, then Blastn was used to identify any sequence with over 90% sequence identity to the ichthyocolid apicoplast SSU. These recovered sequences were then compiled and mapped to the apicoplast SSU tree constructed earlier using RAxML’s evolutionary placement algorithm. The General Time Reversible model with a gamma-distributed rate variation (GTR+CAT) was used along with a minimum bootstrap support threshold of 20% to generate the EPA-placement tree. Those sequences which clustered within the ichthyocolids were considered members of the clade and used to determine overall ichthyocolid prevalence and distribution by cross-referencing them with the original metadata of their respective studies.

### Data and Code Availability

All scripts, organelle assemblies, alignments, and raw tree files used for data analysis can be found on GitHub at https://github.com/delCampoLab/Ichthyocolids. Accession numbers for the sequences used the phylogenetic analyses can be found in Supplementary Table 3. Ichthyocolid apicoplast, mitochondrion, LSU rRNA, and SSU rRNA sequences have been deposited in NCBI GenBank with the following accession numbers: OR826408, OR822199, OR832858, and OR832857 respectively. Raw reads have been deposited in SRA under BioProject PRJNA1041302.

## Supplemental Information Titles & Legends

**Supplementary Table 1. Information on recovered ichthyocolids.**

**Supplementary Table 2. Fish microbiome studies examined for the presence or absence of ichthyocolid reads.**

**Supplementary Table 3. Accession numbers for phylogenetic trees.**

## Notes

https://github.com/delCampoLab/Ichthyocolids

### Competing Interest Statement

The authors have declared no competing interest.

